# Minimizing ion competition boosts volatile metabolome coverage by secondary electrospray ionization orbitrap mass spectrometry

**DOI:** 10.1101/2020.07.27.222505

**Authors:** Jiayi Lan, Jérôme Kaeslin, Giorgia Greter, Renato Zenobi

**Author notes:** Equal contribution (Jiayi Lan, Jérôme Kaeslin).

## Abstract

Secondary electrospray ionization high-resolution mass spectrometry (SESI-HR-MS) is an emerging technique for the detection of volatile metabolites. However, sensitivity and reproducibility of SESI-HRMS have limited its applications in untargeted metabolomics profiling. Ion suppression in the SESI source has been considered to be the main cause. Here, we show that besides ion suppression, ion competition in the C-trap of Orbitrap instruments is another important factor that influences sensitivity and reproducibility of SESI-MS. Instead of acquiring the full mass-to-charge ratio (*m/z*) range, acquisition of consecutive *m/z* windows to minimize the ion competition effect allows the detection of more features. *m/z* window ranges are optimized to fill the C-trap either with an equal number of features or an equal cumulative intensity per window. Considering a balance between maximizing scanning speed and minimizing ion competition, splitting the *m/z* = 50-500 range into 4 windows is selected for measuring human breath and bacterial culture samples on SESI-Orbitrap MS, corresponding to a duty cycle of 2.3 s at a resolution of 140’000. In a small cohort of human subjects, the proposed splitting into 4 windows allows three times more features to be detected compared to the classical full *m/z* range method.

## 1. Introduction

Volatile metabolites provide deep insight into metabolism. Various methods have been developed for the analysis of volatile metabolites and are commonly applied in clinical research, mainly using gas chromatography-mass spectrometry (GC-MS) [1], proton-transfer-reaction mass spectrometry (PTR-MS) [2] and selected-ion flow-tube mass spectrometry (SIFT-MS) [3]. Over the last decade, secondary electrospray ionization (SESI) has emerged and has been widely used to ionize metabolites in the gas phase. Given its modularity [4], SESI can be coupled with various commercial mass spectrometers. Particularly attractive is the coupling to high-resolution mass spectrometers (HRMS), e.g. an Orbitrap. Previous studies have shown that coupling to such a high-resolution instrument offers greater selectivity in feature detection and simplifies metabolite identification.[5, 6] Besides the number of detected features and the resolving power, the reproducibility of the detected features is an important aspect and has been addressed in recent standardization studies.[7, 8]

Although efforts has been made to improve ionization efficiency [9] and feature detection stability [7] of SESI, reproducibility and sensitivity might still be impaired by ion suppression - an effect well known from classical electrospray methods in which a large number of analytes are introduced into the ion source and competition for charge carriers takes place.[10] Besides ion suppression, a large number of analytes are also problematic in other regards: (i) coalescence might results in fewer detected features and reduced mass accuracy[11], (ii) mass accuracy might deteriorate[12], and (iii) impaired sensitivity may lead to many undetected analytes. This reduced sensitivity is sometimes referred to as *ion competition*[13] or *post interface signal suppression*[14] on Orbitraps. This effect arises with matrix-heavy samples when high and low abundant ions compete for space in the capacity-limited C-trap.[14] Previous studies have shown that the ion competition effect can be minimized by applying a strategy called spectral stitching, in which smaller *m/z* windows are acquired and stitched together instead of acquiring the whole *m/z* range in one scan.[15, 16] In the case of an Orbitrap, this avoids the overfilling of the C-trap with the most abundant ions when less abundant ions do not accumulate enough to be detected. However, since multiple scans are required to cover the whole *m/z* range, spectral stitching leads to a significantly longer duty cycle than a full MS scan. Instead of performing spectral stitching with evenly distributed *m/z* windows, it has recently been shown by *Sarvin et al*. in electrospray ionization based flow injection mass spectrometry, that optimized window positions and widths enhance sensitivity while keeping the duty cycle short.[13] In their study, the windows were optimized such that the number of features was equal in every window.

Here, we optimize *m/z* windows to mitigate ion competition on SESI-Orbitrap MS instruments for the real-time monitoring of gaseous analytes. We do this for human breath and bacterial culture headspace, where the monitored event are very time-limited (i.e., an exhalation, which lasts only a few seconds).

## 2. Methods

### 2.1. Real-time SESI-HRMS measurements of human breath and bacterial cultures

The SESI-HRMS system consisted of a commercial SESI source (Super SESI, Fossiliontech, Spain), a flow sensor (Exhalion, Fossiliontech) and an Orbitrap mass spectrometer (Q-Exactive Plus, Thermo Scientific, San Jose, CA). A 20 µm non-coated TaperTip emitter (New Objective, Woburn, MA) was used for electrospray using an aqueous solution of 0.1% formic acid. The sampling line temperature of the SESI source was set to 130 ^*°*^C and the ion chamber temperature was set to 90 ^*°*^C. Breath was directly introduced into the sampling line and the ion chamber of the SESI source through a spirometry filter (MicroGard, Vyaire Medical, USA; 3 cm ID) as previously described [7]. By controlling the exhaust mass flow cutoff of the SESI source at 0.65 L min^*−*1^ and the nitrogen mass flow through the source at 0.35 L min^*−*1^, 0.3 L min^*−*1^ of breath was constantly sampled by the mass spectrometer while subjects were exhaling at a flow rate of 8 L min^1*−*^, similar to the settings of a recent standardization study[7]. For preparing bacteria samples, *Escherichia coli* O9 HS was cultured to an optical density of 0.32, then filtered through a 0.22 *µ*m filter to remove the bacteria. Hemin-supplemented (0.25 mg/*µ*L) brain heart infusion broth was used as medium for culturing bacteria. To sample the bacterial culture’s headspace, 0.3 L min^*−*1^ of humidified nitrogen was flushed through a culture flask containing either medium or culture supernatant, and introduced into the SESI source.

Data was acquired on the Orbitrap in profile mode. For both positive and negative mode acquisition, the following settings were used: an ion transfer tube temperature of 250 ^*°*^C, a resolution of 140’000 at *m/z* = 200, a *m/z* = 50-500 scan range, RF lens set at 55%, and a maximum injection time of 100 ms. The automatic gain control (AGC) target, which regulates the number of ions accumulated in the C-trap and later injects them into the Orbitrap analyzer, was set to 5 · 10^6^. As an instrument blank, data was acquired without introducing human breath or bacterial culture headspace. In parallel to the MS data acquisition, the flow rate and total volume of the gas flows into the SESI source were measured by the Exhalion flow sensor.

Different MS acquisition methods with different scan ranges were used, including a) full scan over a range from *m/z* = 50-500; b) targeted with the narrowest scan window possible from *m/z* = 204.8-205.2; c) twenty-two 20 Da windows with an overlap of 4 Da; d) a variable number *N* of optimized windows covering the *m/z* = 50-500 range splitted either according to the equal-feature number criterion (equal number of robust features per window - the same strategy as in [13]) or the equal-cumulative intensity criterion (equal cumulative intensity per window), with no overlap between windows. Figure 1)i) shows how these ranges are allocated while a subject is exhaling.

**Figure 1:**
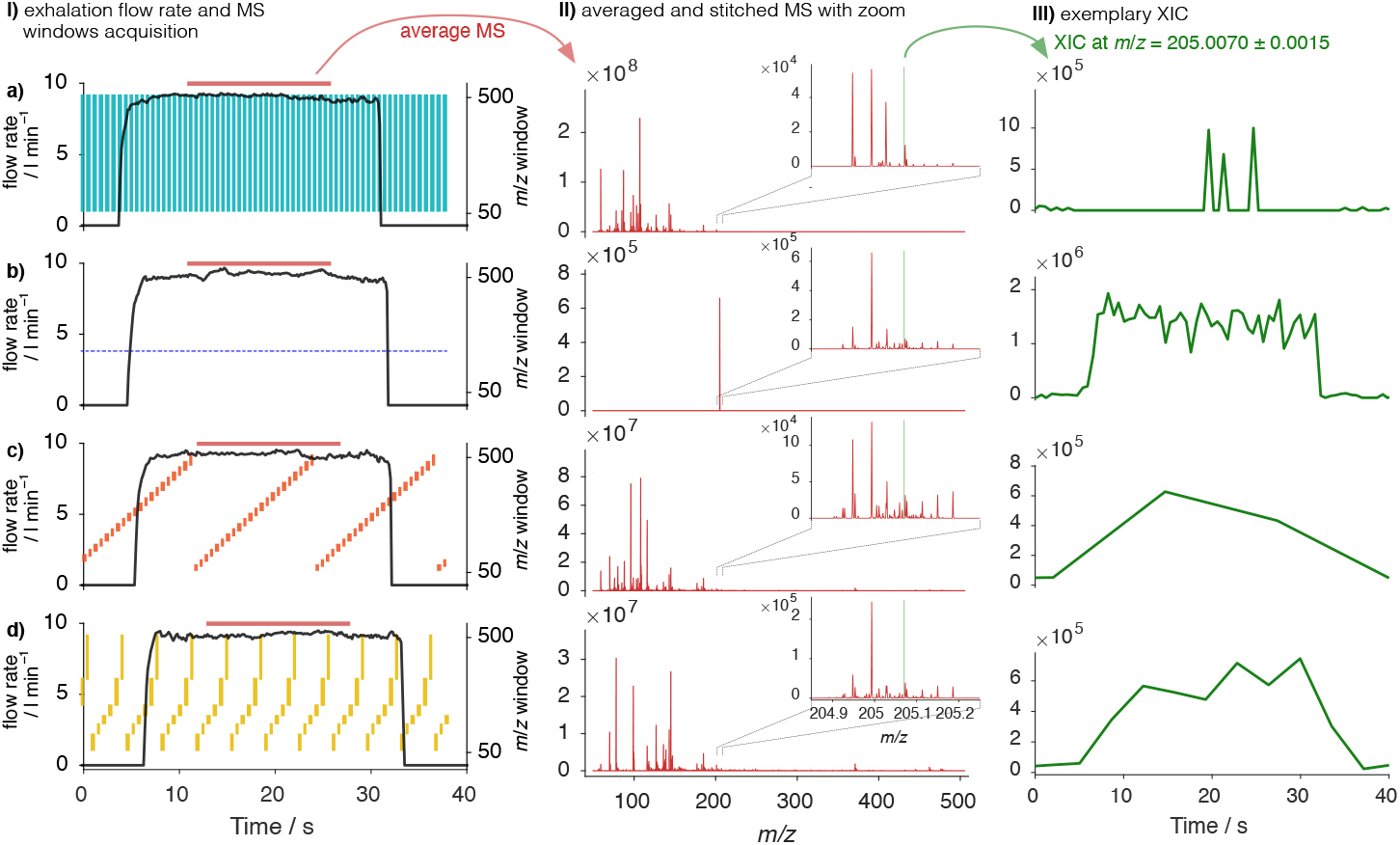
Overview of different acquisition strategies; a) full scan from *m/z* = 50-500; b) targeted from *m/z* = 204.8-205.2; c) 22 sequential 20 Da windows; d) *N* = 6 intensity-optimized windows: i) shows one exhalation of a subject and plots the exhalation flow rate. The scan windows are shown in colors. ii) shows the average spectrum during the exhalation with a zoomed region. iii) shows the extracted ion chromatogram (XIC) of an exemplary selected feature.

### 2.2. Data processing

Raw data generated from the mass spectrometer were first converted into .mzXML format via MSConvert (ProteoWizard [17]). The converted files were then imported into Matlab (version R2019b, MathWorks, Natick, MA) and treated using a Matlab code developed in-house. Briefly, peaks were picked from an overall averaged spectrum of acquired spectra during the experiment. The acquired data was then centroided using the picked peaks, and a .csv file containing the intensities of the peaks per scan was generated for each of the acquired MS datasets.

Statistics and window suggestions were calculated using in-house developed R codes.^1^ For each of the samples, the mean intensity value of the acquired scans was used for quantification. Overlapping regions between two windows were averaged. The peaks were grouped within 5 ppm ranges and the peak intensities were summed as the peak group intensity. The summed peaks were then used to identify robust features using the criteria described by *Sarvin et al*. [13]. *Briefly, a feature was considered “robust” only if it matched the following criteria: i) observed in ≥* 80 % of the biological samples; ii) the median intensity of the biological samples is above 1000 units; iii) signal-to-noise ratio above 4; iv) for breath samples, a relative standard deviation (RSD) across the biological sample injection lower than 30 % and for bacteria culture, lower than 100 %. The windows were then optimized to either contain an equal number of features per window (feature-optimized), or equal cumulative intensity per window (intensity-optimized).

## 3. Results and discussion

### 3.1. Targeted data acquisition method improves sensitivity of SESI-HRMS

In a classical SESI-MS study, full MS scans are acquired repeatedly to track a subject’s exhalation (figure 1a)i)). From this data, an averaged spectrum can be calculated (figure 1a)ii)). Depicting a signal over time gives an idea as to whether the feature is elevated during exhalation or not - as in the case of figure 1a)iii) where the exemplary feature at *m/z* = 205.007 was inconstantly detected by the mass spectrometer. Such features below a certain exhalation-to-background threshold cannot be confidently quantified or attributed to the breath events. Consequently, they are typically filtered out during data analysis.[18]

Based on the hypothesis that ion competition might limit sensitivity in SESI-MS, we investigated how the feature detection performance is influenced by the acquisition method and therefore by the extent that the C-trap is filled with analytes of interest. One exhalation was acquired with a full scan method from *m/z* = 50-500 and with a targeted method from *m/z* = 204.8-205.2 (figure 1a)i) and b)i)) More peaks were observed in the region *m/z* = 204.8-205.2 with the targeted method (figure 1a)ii) vs. b)ii)). Additionally, the signals were clearly elevated during exhalation with the targeted method while the full scan method suffered from a poor exhalation-to-background ratio (exemplary time trace in figure 1a)iii) vs. b)iii)). Figure S1iii) shows that this observation was persistent over multiple exhalations.

The full scan method, which fills the C-trap with all available ions, performs significantly worse than the targeted method, which only traps the ions of interest. This observation supports the hypothesis that ion competition limits sensitivity and robustness of the observed features in full scan SESI-Orbitrap MS. A similar behavior was observed for other targeted mass ranges (*m/z* = 89.8-90.2, 253.9-254.3, 397.9-398.3, 412-412.4, results not shown).

### 3.2. Defining the ion distribution of human breath and bacteria culture

Ion competition effects in untargeted studies can be minimized by spectral stitching.[15][16] We generated a spectral stitching method by using twenty-two 20 Da windows as shown in figure 1 i)c). The stitched spectrum contained significantly more peaks compared to the full scan spectrum (figure 1 ii)a) vs. ii)c)). Moreover, the exemplary *m/z* = 205.007 feature was reliably detected (figure 1 iii)c) and figure S1 iii)c)). Thus, ion competition is mitigated using twenty-two 20 Da windows, however the time resolution is low due to long time to accomplish one duty cycle (*≈* 14 s).

In a more systematic experiment, we compared the detected features by using twenty-two 20 Da windows over the mass range of *m/z* = 50-500 both for human breath and bacteria culture headspace. An overview of the detected features comparing the two methods is given in figure 2. There, the robustly detected features in 22 scans are shown, i.e. comparing one *m/z* = 50-500 full scan with twenty-two stitched 20 Da isolation windows. For breath and bacterial headspace, a 4-to 7-fold increase of robustly detected features is observed. Ion competition particularly suppresses analytes in the higher *m/z* region - presumably because the ion competition is mainly caused by high abundant low mass matrix compounds. In fact, the ion competition-reduced feature distribution more closely resembles the distribution reported on a SESI-quadrupole time-of-flight (QTOF) instrument[19], where ion competition is not expected because of the ion-trap free nature of a QTOF instrument.

**Figure 2:**
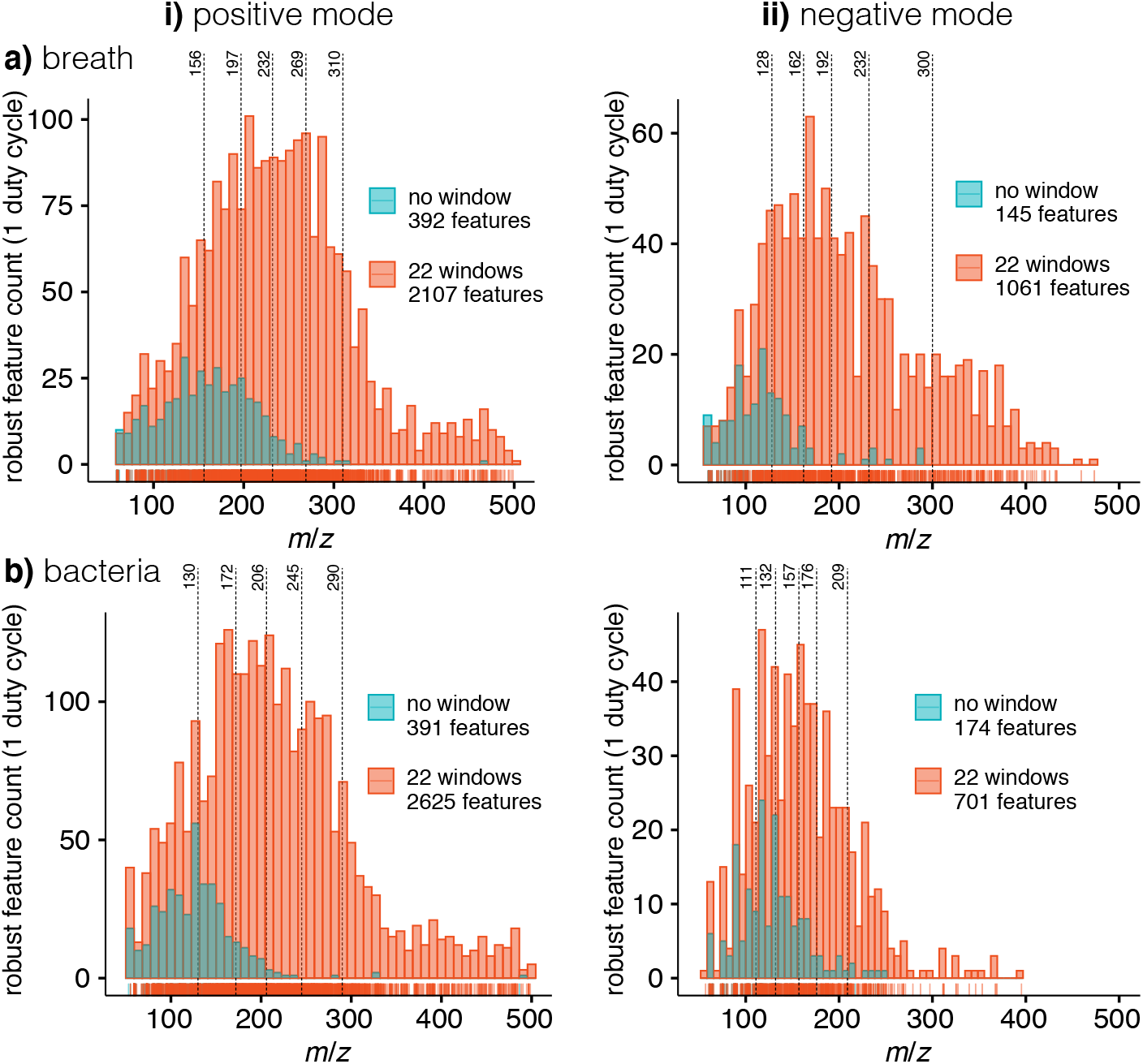
Feature distribution for different acquisition methods with one duty cycle (one *m/z* = 50-500 scan vs. twenty-two stitched 20 Da windows): shows the feature distribution in breath (a) and bacterium culture headspace (b) observed with no *m/z* windows (i.e. full scan *m/z* = 50-500) in blue. Using twenty-two 20 Da windows, more features are detected (in red) due to minimized ion competition. The feature distribution is shown for positive i) and negative ii) polarity.

While the twenty-two windows method significantly enhances the number of observed features, it also prolongs the duty cycle. To account for the longer duty cycle of the twenty-two windows method, the two strategies can be benchmarked by comparing them, with respect to an identical acquisition time (22 scans, roughly 14 s). This comparison takes into account that the *m/z* = 50-500 full scan method has a better counting statistic, i.e. a potential feature is scanned 22 times more in the same period of time. Nonetheless, figure S2 shows that the twenty-two windows strategy outperforms the no windows strategy by a factor of two in terms of detected robust features for positive mode in breath and bacteria. The same holds true for the negative mode in breath, but in bacteria where both methods perform similarly well. Overall, the performance of the two methods within an identical acquisition time is comparable in the low mass region. However, the twenty-two windows method clearly performs better in the mid-to high mass region.

### 3.3. Optimized windows for human breath and bacteria headspace acquisition

Assuming that the twenty-two windows feature distribution in figure 2 represents the ion-competition free distribution, the optimized windows’ center and widths can be determined by allocating an equal number of robust features into one optimized window as outlined before in [13]. The borders of these “feature optimized” windows are marked with a dashed line in figure 2 for *N* = 6 windows. Additionally, we hypothesized that ion competition is more efficiently controlled by allocating the windows not to contain an equal number of robust features but instead to contain an equal cumulative intensity. This is because ion competition is not driven by the robust features themselves but rather by highly abundant ions - independent from whether they arise from the sample or from the instrument. Consequently, we also calculated “intensity-optimized” windows to containing an equal amount of cumulative intensity per window during exhalation or headspace acquisition as shown in figure S3.

2 to 16 optimized windows were calculated based on the “feature-optimized” and the “intensity-optimized” criteria. The corresponding window limits can be found in table S1 for breath and S2 for bacterial headspace, respectively. It should be mentioned that these suggested windows are dependent on the matrix and the general background level of ions arising from the instrument and environment. However, we assume that the matrix of human breath has an inter-subject similarity. Moreover, the window splitting is expected to mitigate ion competition even if the exact window breaking point might not be exactly ideal. Thus, the reported window limits could directly be adapted by other groups. An example for 6 intensity-optimized windows is shown in figure 1d) and S1d). The exemplary feature *m/z* = 205.007 is robustly detected. Therefore, the advantages of window optimization are that it allows the detection of suppressed features (figure 1a) vs. d)), is untargeted, (figure 1b) vs. d)) and has an acceptable time resolution (figure 1c) vs. d), time for one duty cycle reduced from *≈* 14 s for 22 windows to *≈* 3 s s for 6 windows).

We reanalyzed human breath and bacterial headspace with *N* = 2, 4, 6, 10 feature- and intensity-optimized windows. The robustly detected features for these window settings are benchmarked with our previously detected robust features of the *m/z* = 50-500 full scan and the twenty-two 20 Da windows strategy in figure 3 using an equal acquisition time. Multiple conclusions can be drawn from this comparison: first, the window splitting always outperforms the *m/z* = 50-500 full scan method. Second, 4 or 6 windows seem to be a good compromise between minimized ion competition and satisfying counting statistics. Third, the intensity-optimized windows have a tendency of outperforming the feature optimized windows, except in the bacterial headspace where the opposite behavior is observed. This supports our hypothesis that windows should preferentially be optimized to contain equal cumulative intensity per window since the sum of charges ultimately drives ion competition.

**Figure 3:**
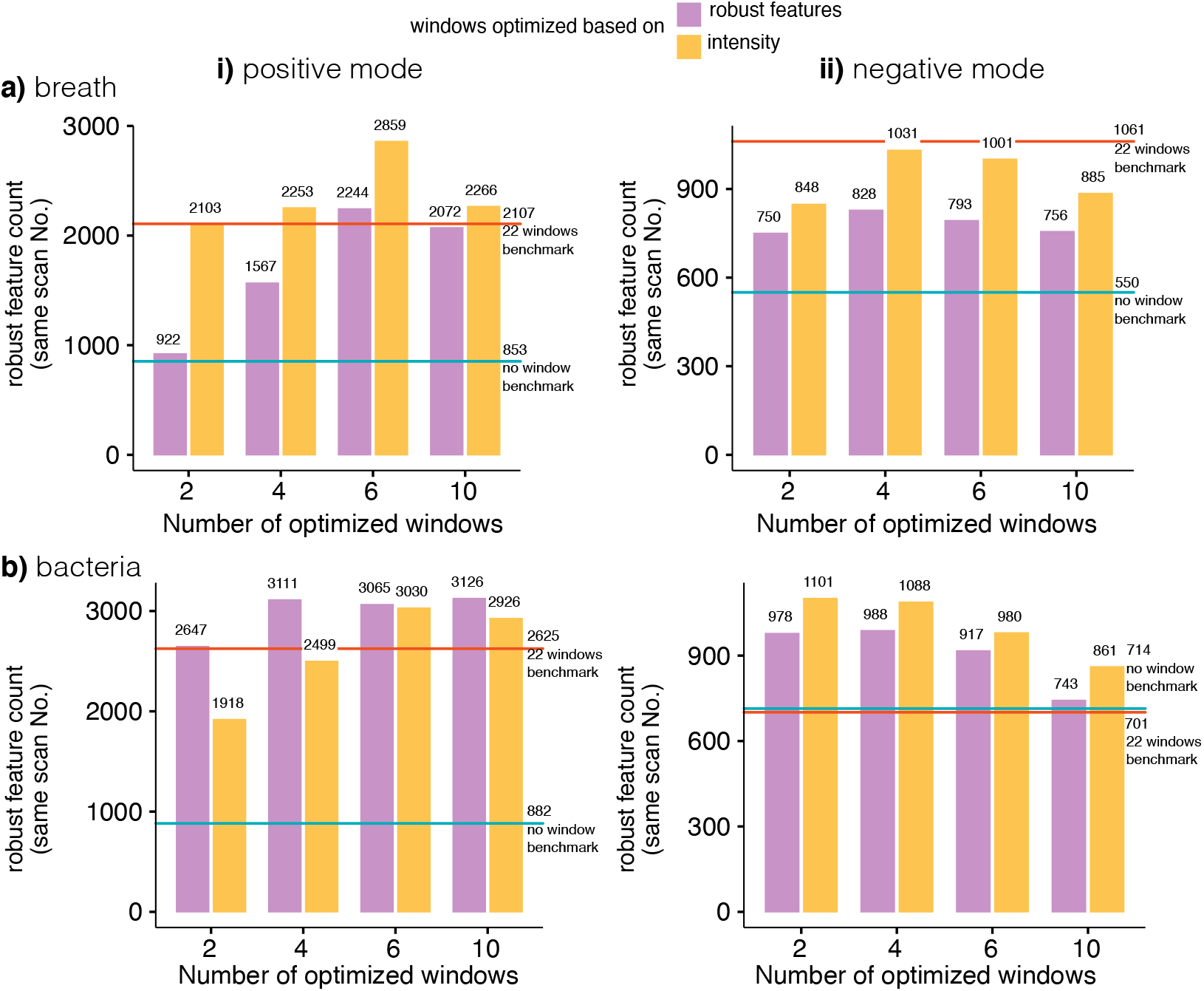
Numbers of robust features detected using *N* = 2, 4, 6, 10 windows in (a) breath and in (b) bacterium culture headspace in 20 scans (*≈* 11 s of scanning time). Both feature-optimized (purple) and intensity-optimized (orange) windows were tested. Moreover, benchmarks from the no windows *m/z* = 50-500 full scans and twenty-two 20 Da windows are represented as a line. Both polarities are shown.

*N* = 4 intensity-optimized windows seem to be the best compromise between reduced ion competition, counting statistics and time resolution. This corresponds to *≈* 2.3 s to scan through the *m/z* = 50-500 range once at a resolution of 140’000 on the Orbitrap Q-Exactive Plus.

### 3.4. Validating the proposed optimized windows in a larger cohort

The human breath windows optimization process from the previous sections is based on only one subject. However, the applicability of these optimized windows may be dependent on inter-subject variability of the breath matrix. We hypothesize that the breath matrix is comparable among different subjects. To support this assumption, we assessed our suggested optimized windows with a cohort of seven people, mimicking the setting of a small scale clinical study. In both positive and negative mode, the median number of robust features detected in individual subjects significantly increased by using 4 intensity-optimized windows, compared to the classical no window approach figure 4a). Moreover, consistency of detected features among subject is also critical for data completeness and statistics. Defining a feature to be consistent if detected in at least 80% of the subjects (*≥* 6 subjects), 455 consistent features were detected using 4 intensity-optimized windows in positive mode and 760 in negative mode. In both cases, the consistent features are significantly increased compared to 196 (positive mode) and 212 (negative mode) obtained using the full scan approach (figure 4b) and c)).

**Figure 4:**
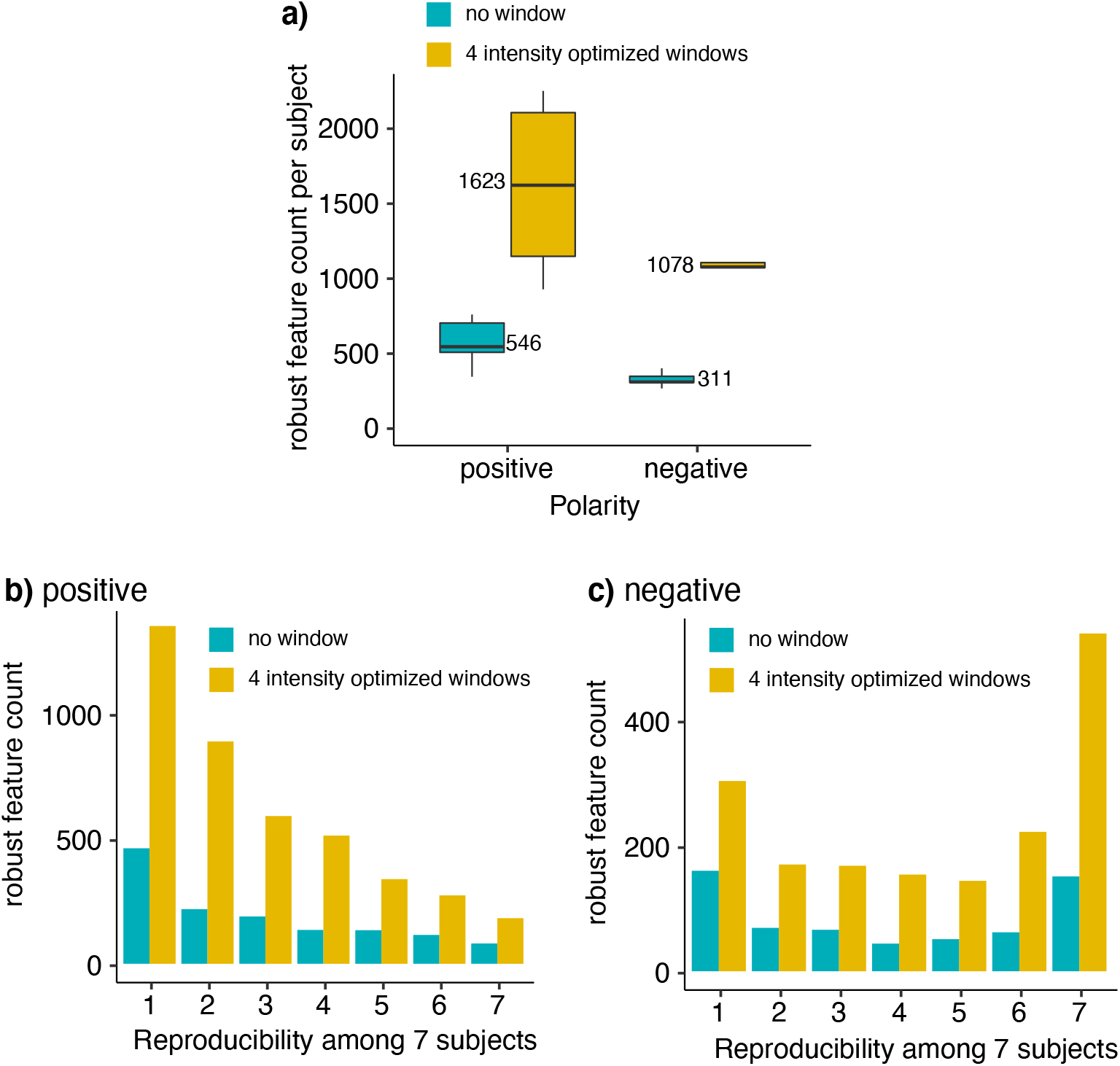
a) Robust features detected per subject. Reproducibility of the detected features among seven subjects in b) positive and c) negative mode.

To assess the quantification stability of the intensity-optimized 4 windows method, an intensity ratio was calculated between 4 feature pairs for each of the replicates. The coefficient of variation within a subject was used as an indication of the quantification stability, assuming the ratio should be stable for one subject over all the replicates (figure 5). When features are robustly detected in both methods, variations are expected to be comparable, independent of whether the pair of features was measured in the same scan window (positive: pyridine/isoprene) or in different scan windows (positive: phenylalanine/isoprene, negative: valeric acid/*m/z* = *−*89.023). For these features, the intensity ratio’s coefficient of variation was comparable between the two methods, indicating that they perform similarly in terms of quantification. For one of the feature pairs in different windows in negative mode (2-hydroxyisobutyric acid/*m/z* = *−*89.023), the coefficient of variation was lower using 4 intensity-optimized windows. This is due to the fact that an analyte signal is underestimated if ion competition takes place [20]. This can be seen through figure S6, the signal variation of 2-hydroxyisobutyric acid is much higher when using the no window approach comparing to the intensity-optimized 4 windows method.

**Figure 5:**
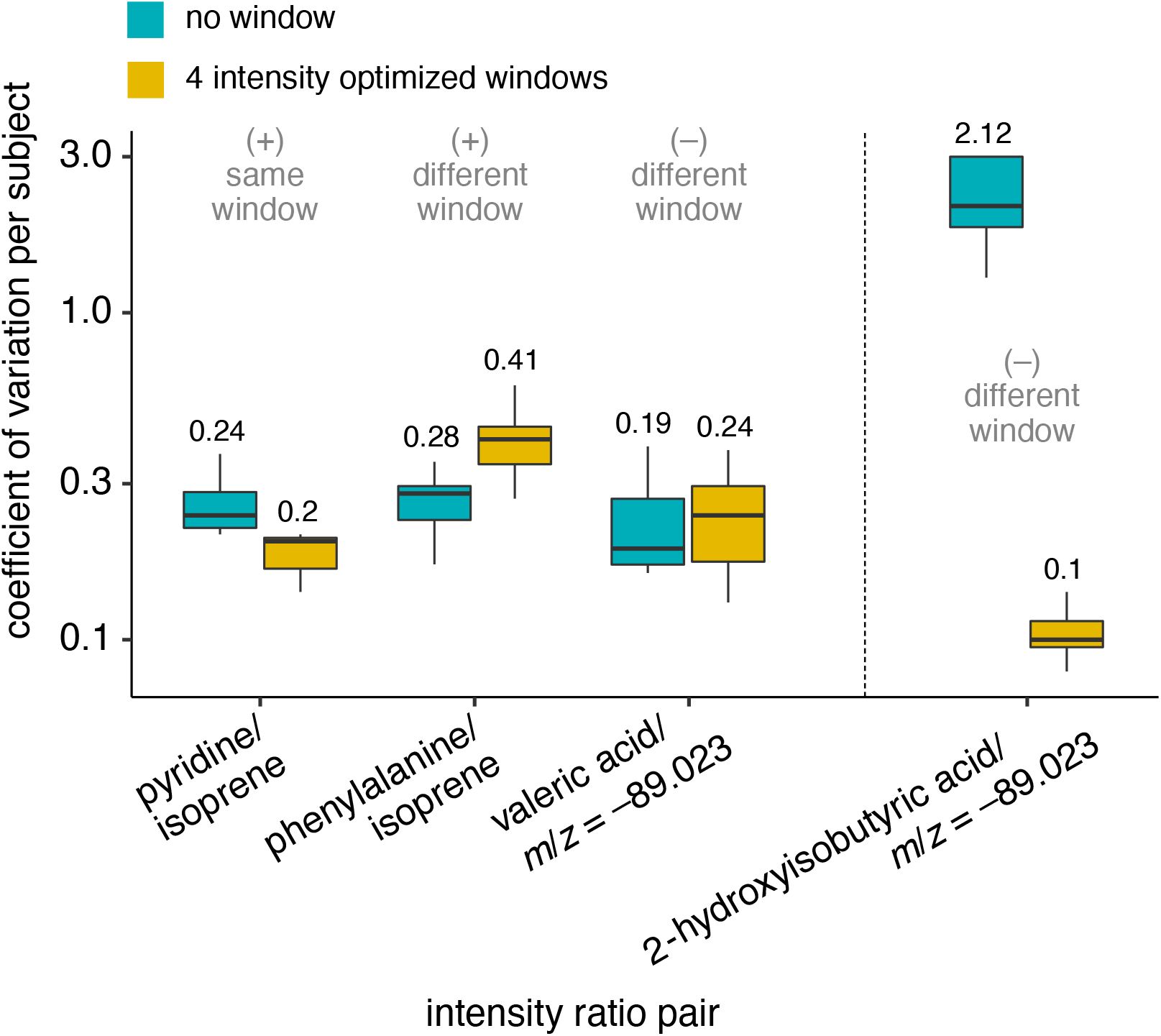
Coefficient of variation of the intensity ratio per subject between 4 features pairs. The labels state the polarities of the features and whether they are located in the same or a different window.

Sensitivity was also assessed by plotting the distribution of signal-to-noise ratios of features detected in both the full scan method and the 4 intensity-optimized window method (figure S5). Only features that have a noise intensity above zero have been considered. Both methods show similar distributions, suggesting the sensitivity of the two methods are at the same level as we expected, since ion suppression hampers both the detection and quantification of very low abundant features while high abundant features are not affected. Linear detection range of the two SESI-MS methods was not compared directly, due to complexity of preparing standards mimicking breath (humid VOC/aerosol mixture). We believe the post-ionization behaviour would be identical to what has been described for electrospray ionization (ESI).([15]), that the linear dynamic range was extended by preventing an overfilling of the C-trap. In summary, the quantification capabilities of the detected features seems to be comparable or even better using the intensity-optimized 4 windows method compared to *m/z* = 50-500 full scan method.

## 4. Conclusions

We show that ion competition indeed limits the sensitivity and and robustness of SESI-Orbitrap MS for analyzing volatiles. For targeted applications, our findings show that a narrow *m/z* window is superior to an untargeted full MS acquisition. For untargeted studies, using spectral stitching with 20 Da windows for acquisition enhances the sensitivity and robustness but limits the sampling frequency due to the prolonged duty cycle. By optimizing the *m/z* window’s positions and widths, a compromise between sensitivity and duty cycle can be found. We demonstrate this approach by using 4 optimized *m/z* windows for the analysis of volatiles in breath and bacteria culture. A significant increase in sensitivity and robustness is observed while the duty cycle is still acceptable. Moreover, different sets of optimized windows for the analysis of breath or bacteria culture volatiles are proposed based on the feature number distribution or the cumulative intensity distribution.

## Acknowledgments

This work is part of the Zurich Exhalomics flagship project under the umbrella of “University Medicine Zurich*/*Hochschulmedizin Zürich”.

## Supplementary figures

**Figure S1:**
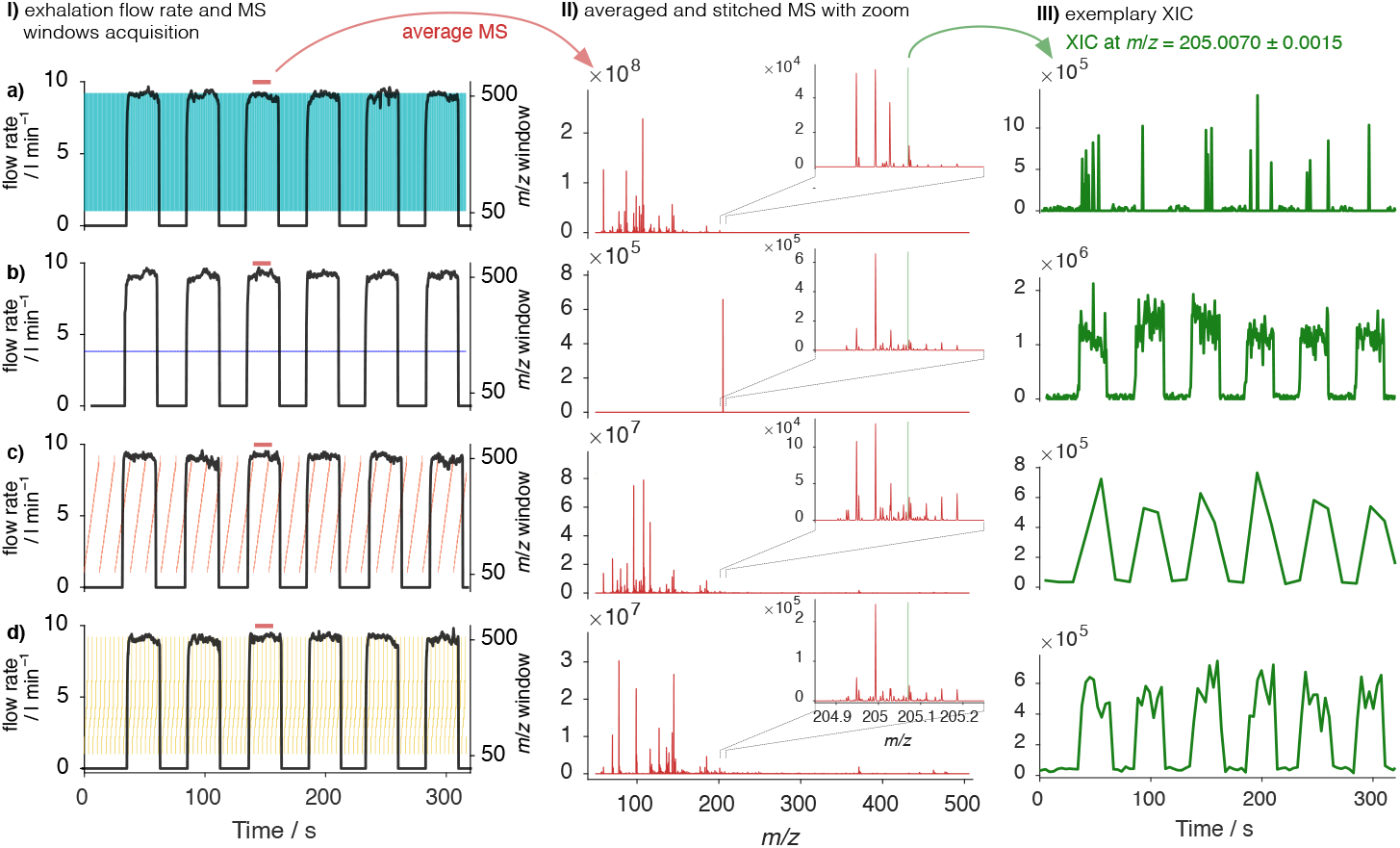
Overview of different acquisition strategies over multiple exhalations; a) full scan from *m/z* = 50-500; b) targeted from *m/z* = 204.8-205.2; c) 22 sequential 20 Da windows; d) *N* = 6 intensity-optimized windows: i) shows multiple exhalations of a subject and plots the exhalation flow rate. The scan windows are shown in colors. ii) Shows the average spectrum during the exhalation with a zoomed region. iii) shows the extracted ion chromatogram (XIC) of an exemplary selected feature.

**Figure S2:**
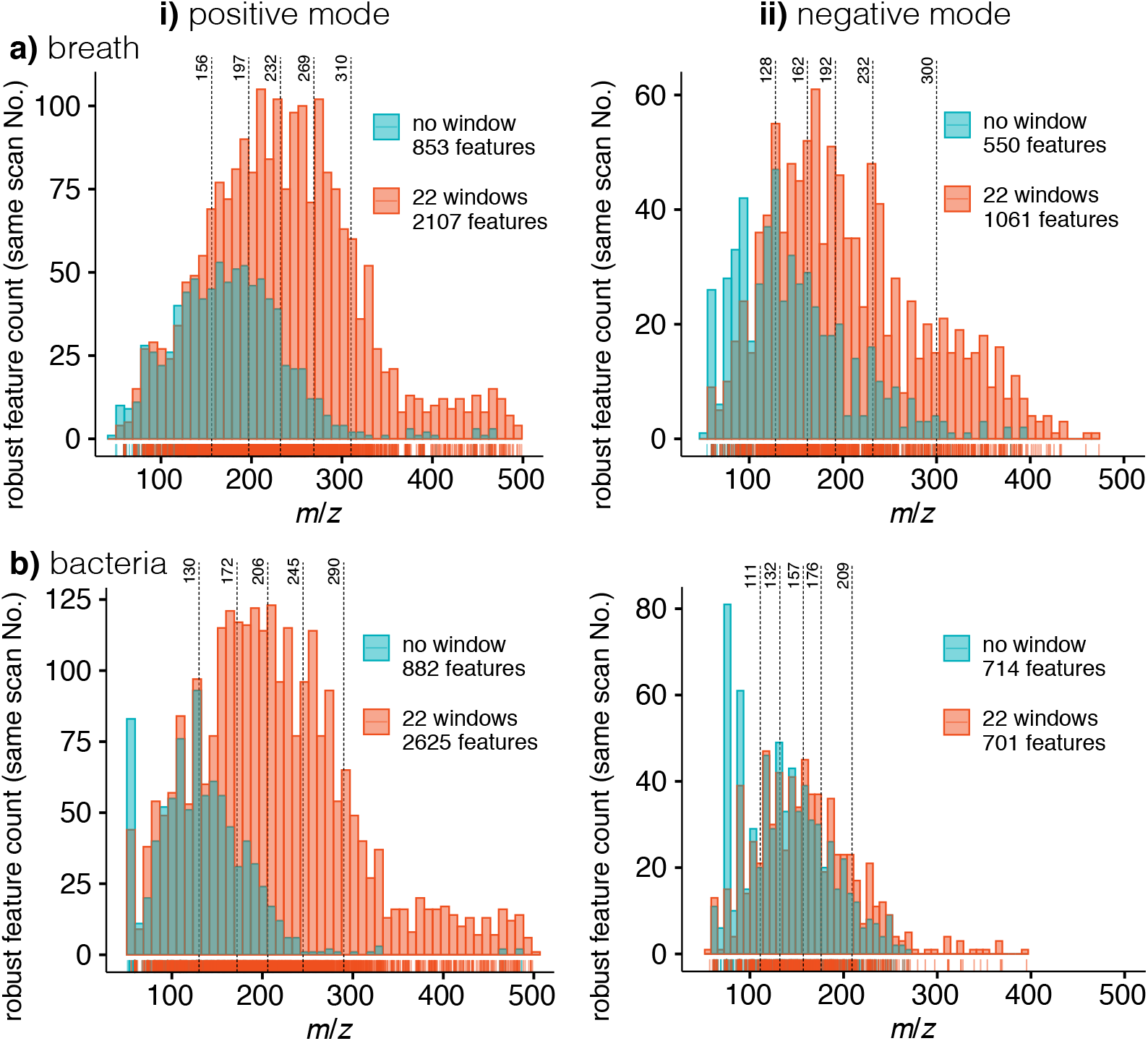
Feature distribution for different acquisition methods comparing an identical acquisition time (twenty-two *m/z* = 50-500 scans vs. twenty-two stitched 20 Da windows): shows the feature distribution in breath (a) and bacterium culture headspace (b) observed with no *m/z* windows (i.e. full scan *m/z* = 50-500) in blue. Using twenty-two 20 Da windows, more features are detected (in red) due to minimized ion competition. The feature distribution is shown for positive i) and negative ii) polarity.

**Figure S3:**
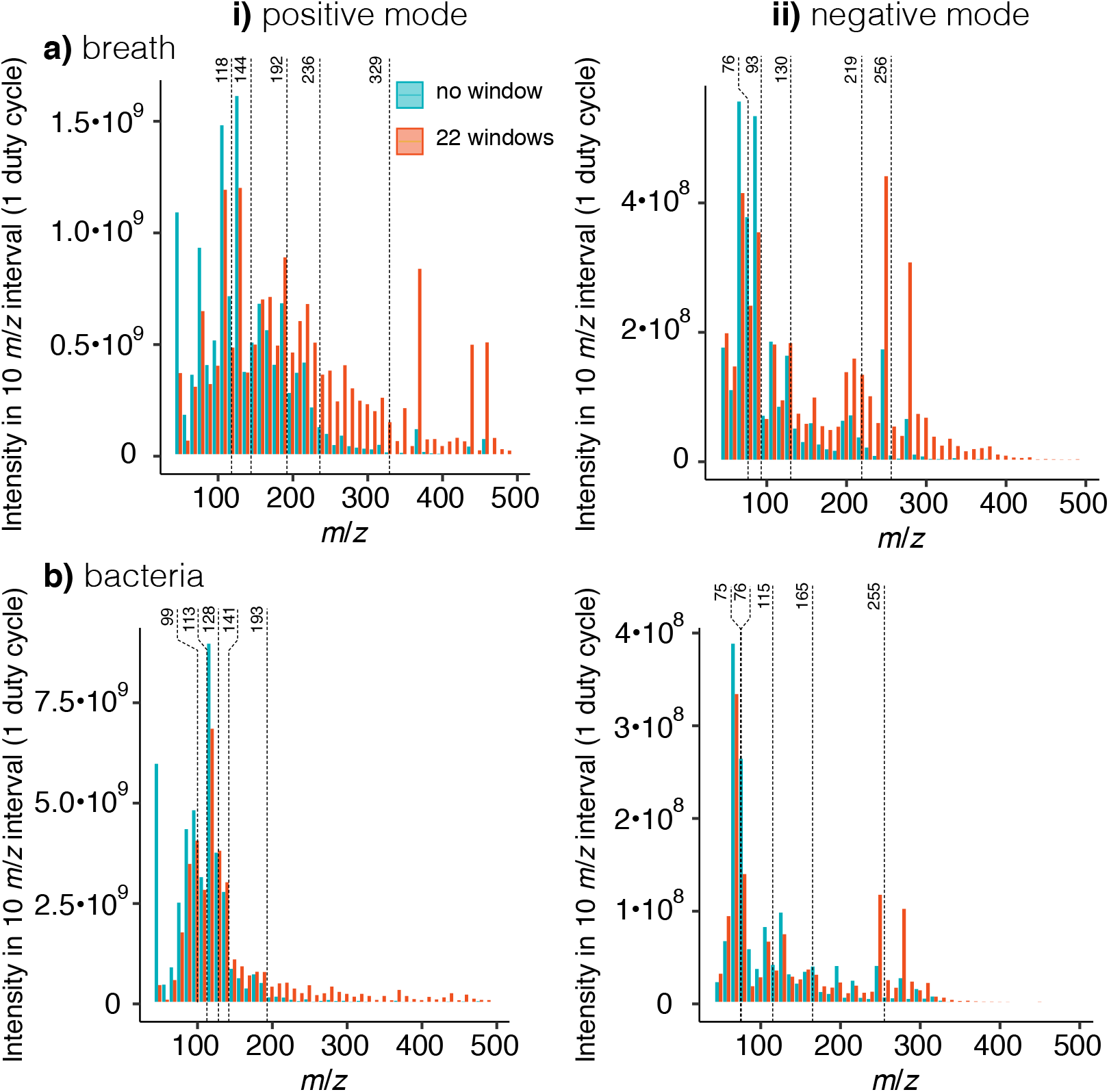
Intensity distribution for different acquisition methods with one duty cycle (one *m/z* = 50-500 scan vs. twenty-two stitched 20 Da windows) in a) breath and in b) bacterium culture headspace. The intensity distribution is shown for positive i) and negative ii) polarity.

**Figure S4:**
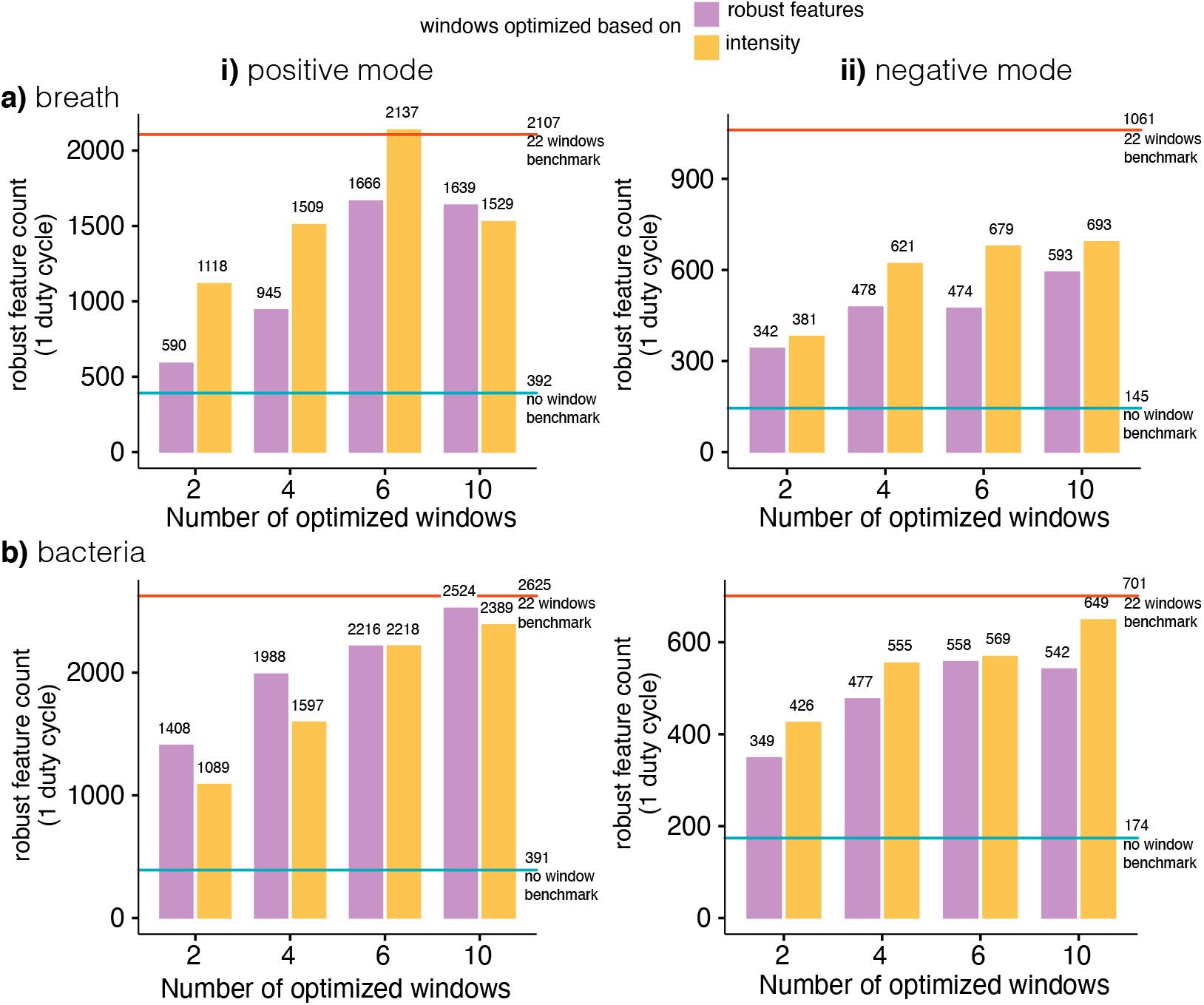
Numbers of robust features detected using *N* = 2, 4, 6, 10 windows in (a) breath and in (b) bacterium culture headspace in one duty cycle. Both feature-optimized (purple) and intensity-optimized (orange) windows were tested. Moreover, benchmarks from the no windows *m/z* = 50-500 full scans and twenty-two 20 Da windows are represented as a line. Both polarities are shown.

**Figure S5:**
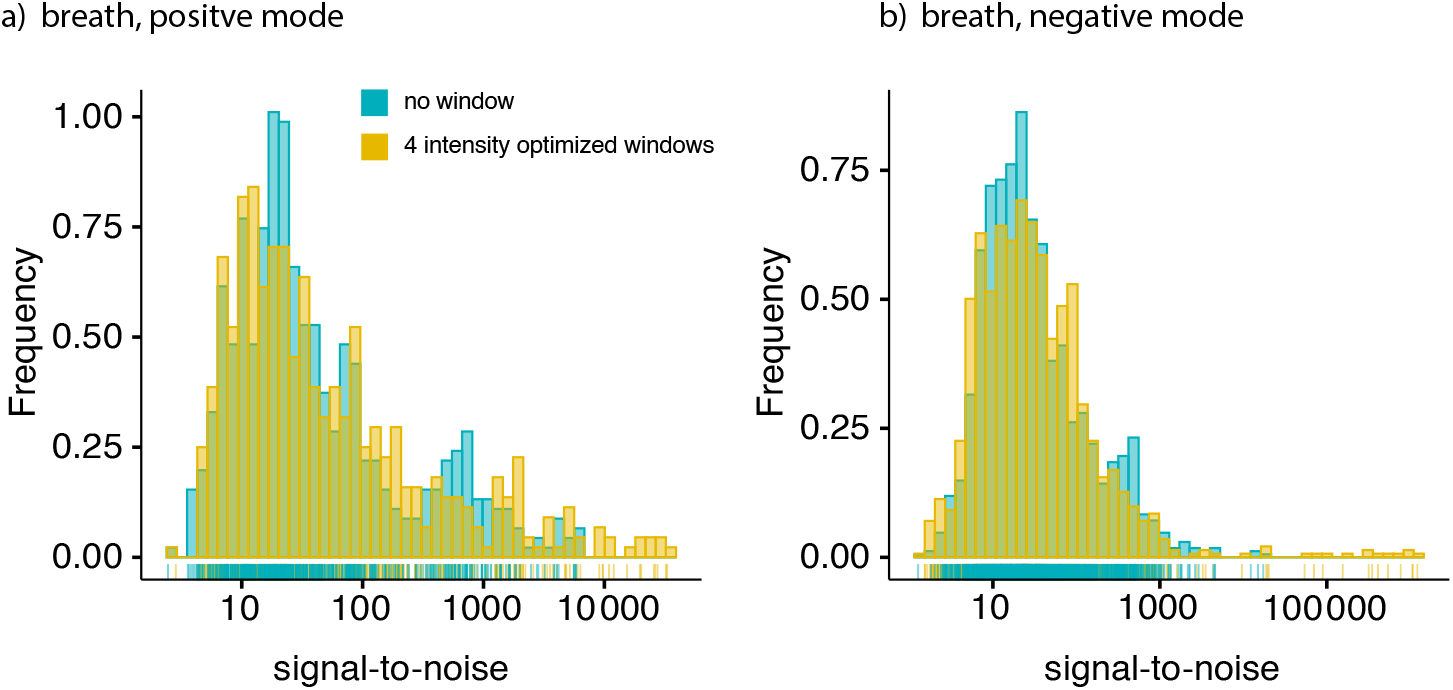
Signal-to-noise ratio distribution of features robustly detected in both methods for positive mode and b) negative polarity.

**Figure S6:**
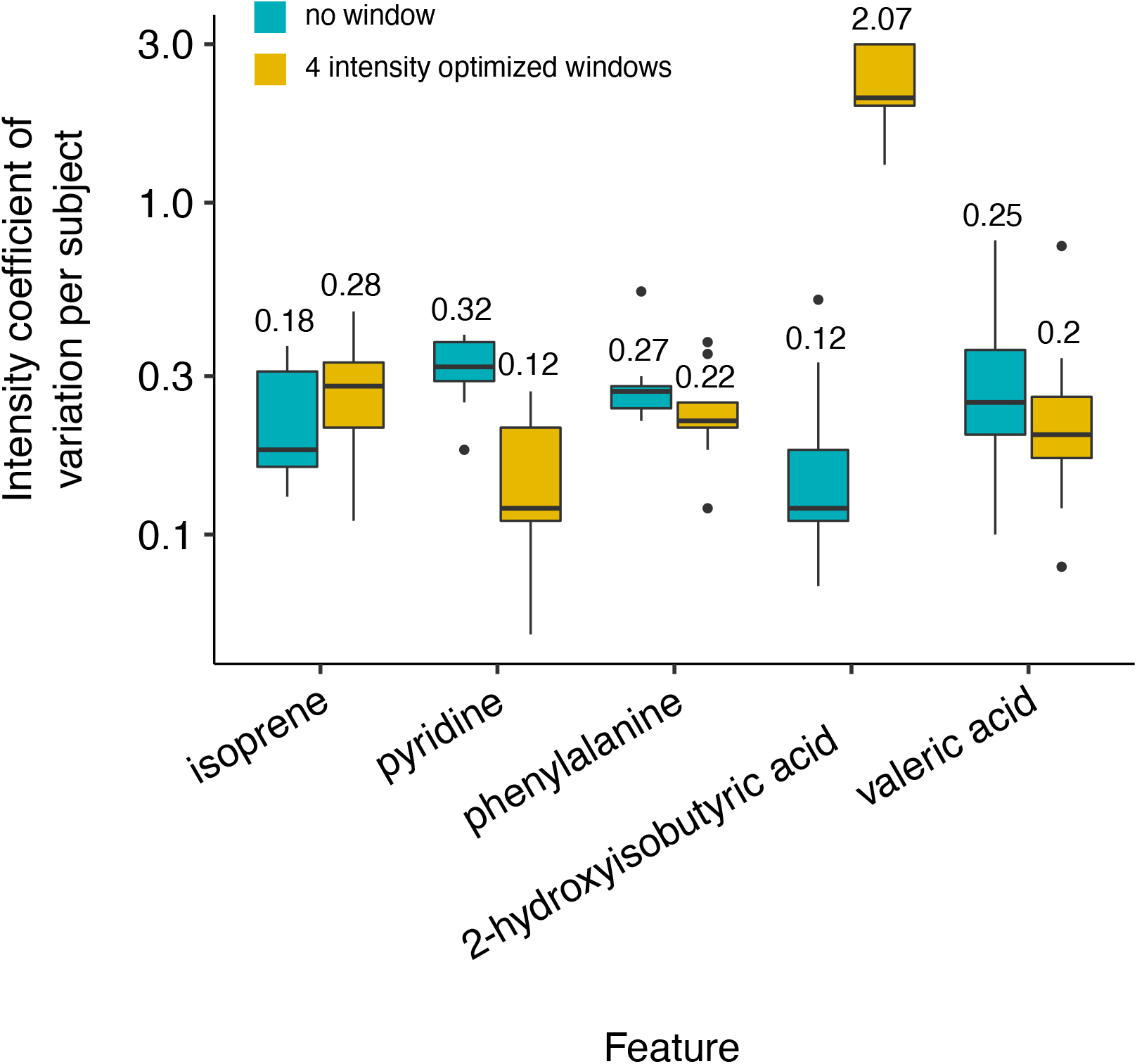
Coefficient of variation of the intensity per subject of 5 features. The features were used to calculate the ratio variation in figure 5.

**Table S1:** Optimized *m/z* windows in human breath. The windows were optimized to either contain an equal number of features or an equal cummulative intensity. Different numbers *N* of windows are proposed such that an analyst can compromise between sensitivity and duty cycle.

| $N$ | 1 | 2 | 3 | 4 | 5 | 6 | 7 | 8 | 9 | 10 | 11 | 12 | 13 | 14 | 15 | 16 |
| --- | --- | --- | --- | --- | --- | --- | --- | --- | --- | --- | --- | --- | --- | --- | --- | --- |
| + breath feature optimized |  |  |  |  |  |  |  |  |  |  |  |  |  |  |  |  |
| 2 | 50-232 | 232-500 |  |  |  |  |  |  |  |  |  |  |  |  |  |  |
| 4 | 50-179 | 179-232 | 232-288 | 288-500 |  |  |  |  |  |  |  |  |  |  |  |  |
| 6 | 50-156 | 156-197 | 197-232 | 232-269 | 269-310 | 310-500 |  |  |  |  |  |  |  |  |  |  |
| 8 | 50-144 | 144-179 | 179-207 | 207-232 | 232-259 | 259-288 | 288-327 | 327-500 |  |  |  |  |  |  |  |  |
| 10 | 50-133 | 133-166 | 166-190 | 190-211 | 211-232 | 232-254 | 254-274 | 274-299 | 299-339 | 339-500 |  |  |  |  |  |  |
| 12 | 50-127 | 127-156 | 156-179 | 179-197 | 197-214 | 214-232 | 232-251 | 251-269 | 269-288 | 288-310 | 310-353 | 353-500 |  |  |  |  |
| 14 | 50-121 | 121-150 | 150-169 | 169-187 | 187-203 | 203-217 | 217-232 | 232-248 | 248-262 | 262-278 | 278-294 | 294-317 | 317-363 | 363-500 |  |  |
| 16 | 50-116 | 116-144 | 144-163 | 163-179 | 179-193 | 193-207 | 207-219 | 219-232 | 232-246 | 246-259 | 259-272 | 272-288 | 288-303 | 303-327 | 327-376 | 376-500 |
| + breath intensity optimized |  |  |  |  |  |  |  |  |  |  |  |  |  |  |  |  |
| 2 | 50-195 | 195-500 |  |  |  |  |  |  |  |  |  |  |  |  |  |  |
| 4 | 50-132 | 132-195 | 195-277 | 277-500 |  |  |  |  |  |  |  |  |  |  |  |  |
| 6 | 50-118 | 118-144 | 144-192 | 192-236 | 236-329 | 329-500 |  |  |  |  |  |  |  |  |  |  |
| 8 | 50-109 | 109-132 | 132-161 | 161-193 | 193-224 | 224-277 | 277-371 | 371-500 |  |  |  |  |  |  |  |  |
| 10 | 50-98 | 98-119 | 119-137 | 137-167 | 167-193 | 193-219 | 219-249 | 249-302 | 302-372 | 372-500 |  |  |  |  |  |  |
| 12 | 50-88 | 88-118 | 118-130 | 130-144 | 144-169 | 169-191 | 191-211 | 211-235 | 235-272 | 272-327 | 327-383 | 383-500 |  |  |  |  |
| 14 | 50-84 | 84-115 | 115-122 | 122-136 | 136-157 | 157-177 | 177-193 | 193-211 | 211-228 | 228-257 | 257-297 | 297-358 | 358-429 | 429-500 |  |  |
| 16 | 50-80 | 80-101 | 101-118 | 118-123 | 123-134 | 134-153 | 153-170 | 170-185 | 185-199 | 199-219 | 219-235 | 235-261 | 261-299 | 299-368 | 368-431 | 431-500 |
| - breath feature optimized |  |  |  |  |  |  |  |  |  |  |  |  |  |  |  |  |
| 2 | 50-192 | 192-500 |  |  |  |  |  |  |  |  |  |  |  |  |  |  |
| 4 | 50-145 | 145-192 | 192-256 | 256-500 |  |  |  |  |  |  |  |  |  |  |  |  |
| 6 | 50-128 | 128-162 | 162-192 | 192-232 | 232-300 | 300-500 |  |  |  |  |  |  |  |  |  |  |
| 8 | 50-119 | 119-145 | 145-168 | 168-192 | 192-224 | 224-256 | 256-322 | 322-500 |  |  |  |  |  |  |  |  |
| 10 | 50-113 | 113-135 | 135-154 | 154-172 | 172-192 | 192-214 | 214-240 | 240-282 | 282-334 | 334-500 |  |  |  |  |  |  |
| 12 | 50-110 | 110-128 | 128-145 | 145-162 | 162-174 | 174-192 | 192-211 | 211-232 | 232-256 | 256-300 | 300-342 | 342-500 |  |  |  |  |
| 14 | 50-103 | 103-124 | 124-138 | 138-152 | 152-166 | 166-178 | 178-192 | 192-208 | 208-228 | 228-244 | 244-274 | 274-312 | 312-352 | 352-500 |  |  |
| 16 | 50-100 | 100-119 | 119-132 | 132-145 | 145-158 | 158-168 | 168-180 | 180-192 | 192-204 | 204-224 | 224-236 | 236-256 | 256-289 | 289-322 | 322-356 | 356-500 |
| - breath intensity optimized |  |  |  |  |  |  |  |  |  |  |  |  |  |  |  |  |
| 2 | 50-161 | 161-500 |  |  |  |  |  |  |  |  |  |  |  |  |  |  |
| 4 | 50-90 | 90-161 | 161-255 | 255-500 |  |  |  |  |  |  |  |  |  |  |  |  |
| 6 | 50-76 | 76-93 | 93-130 | 130-219 | 219-256 | 256-500 |  |  |  |  |  |  |  |  |  |  |
| 8 | 50-75 | 75-89 | 89-96 | 96-141 | 141-215 | 215-255 | 255-283 | 283-500 |  |  |  |  |  |  |  |  |
| 10 | 50-75 | 75-86 | 86-93 | 93-114 | 114-138 | 138-205 | 205-233 | 233-255 | 255-283 | 283-500 |  |  |  |  |  |  |
| 12 | 50-61 | 61-76 | 76-89 | 89-93 | 93-105 | 105-130 | 130-165 | 165-219 | 219-247 | 247-256 | 256-284 | 284-500 |  |  |  |  |
| 14 | 50-60 | 60-75 | 75-76 | 76-89 | 89-93 | 93-97 | 97-129 | 129-159 | 159-205 | 205-222 | 222-255 | 255-256 | 256-284 | 284-500 |  |  |
| 16 | 50-60 | 60-75 | 75-76 | 76-89 | 89-93 | 93-94 | 94-117 | 117-132 | 132-171 | 171-207 | 207-227 | 227-255 | 255-256 | 256-283 | 283-297 | 297-500 |

**Table S2:**
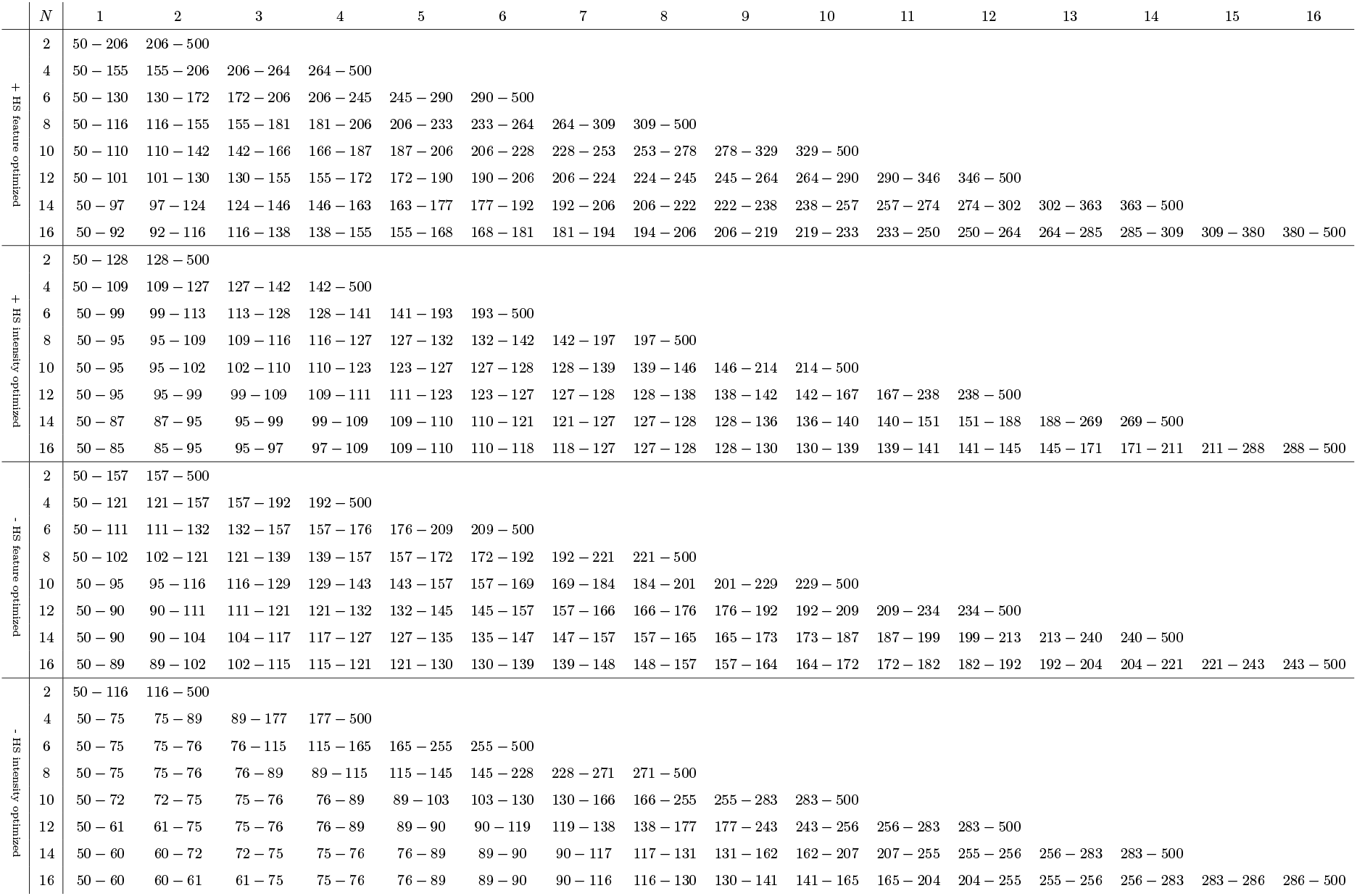
Optimized *m/z* windows in bacteria culture headspace. The windows were optimized to either contain an equal number of features or an equal cummulative intensity. Different numbers *N* of windows are proposed such that an analyst can compromise between sensitivity and duty cycle.

## Footnotes

1 The R codes and the raw data can be found under https://doi.org/10.3929/ethz-b-000445560

## Notes

### Competing Interest Statement

The authors have declared no competing interest.

